# Citric Acid Water as an Alternative to Water Restriction for High-Yield Mouse Behavior

**DOI:** 10.1101/2020.03.02.973016

**Authors:** Anne E Urai, Valeria Aguillon-Rodriguez, Inês C Laranjeira, Fanny Cazettes, The International Brain Laboratory, Zachary F Mainen, Anne K Churchland

## Abstract

Powerful neural measurement and perturbation tools have positioned mice as an ideal species for probing the neural circuit mechanisms of cognition. Crucial to this success is the ability to motivate animals to perform specific behaviors. One successful strategy is to restrict their water intake, rewarding them with water during a behavioral task. However, water restriction requires rigorous monitoring of animals’ health and hydration status and can be challenging for some mice.

We present an alternative that allows mice more control over their water intake: free home-cage access to water, made slightly sour by a small amount of citric acid (CA). In a previous study, rats with free access to CA water readily performed a behavioral task for water rewards, although completing fewer trials than under water restriction (Reinagel, 2018). We here extend this approach to mice and confirm its robustness across multiple laboratories.

Mice reduced their intake of CA water while maintaining healthy weights. Continuous home-cage access to CA water only subtly impacted their willingness to perform a decision-making task, in which they were rewarded with sweetened water. When free CA water was used instead of water restriction only on weekends, learning and decision-making behavior were unaffected. CA water is thus a promising alternative to water restriction, allowing animals more control over their water intake without interfering with behavioral performance.

**Significance Statement:** High-throughput, reliable behavioral training is a key requirement for the use of mice in behavioral and systems neuroscience, but depends crucially on ability to motivate animals to perform specific behaviors. Here, we present an alternative method to commonly used methods of water restriction: free home-cage access to water, made slightly sour by a small amount of citric acid. This non-labor-intensive, low-error option benefits animal health without hindering behavioral training progress. Citric acid water can serve as a reliable and standardized strategy to achieve high quality task behavior, further facilitating the use of mice in high-throughput behavioral studies.

## Introduction

The mouse is an indispensable species for systems neuroscience, thanks to a rich set of available tools to record and manipulate brain structure and function, combined with the knowledge of teaching mice specific behavioral tasks. While long thought to be beyond the capacities of the mouse (Abbott, 2010), mice are now routinely trained to perform abstract sensory, navigational and decision-making tasks (Carandini and Churchland, 2013; Guo et al., 2014; Goltstein et al., 2018). Crucial to this success is the ability to motivate them to perform specific behaviors.

One successful strategy is to restrict animals’ water access, and reward them with fluids for performing behavioral tasks (Skinner, 1936; Toth and Gardiner, 2000; Guo et al., 2014; Goltstein et al., 2018). The mouse’s ability to thrive on little water matches its evolutionary past on steppes and other dry environments (Fertig and Edmonds, 1969). In a laboratory setting, however, water restriction requires rigorous monitoring of animal’s health and hydration status, usually with daily weighing and precisely measured water intake (Toth and Gardiner, 2000).

We here consider a complementary, alternative approach: free access to water, in which a small amount of citric acid (CA) has been dissolved. CA is a food preservative which makes water taste slightly sour. Rats reduce their intake of CA water without getting sick or dehydrated (Watson et al., 1986). Moreover, free CA water has only subtle impacts on rats’ willingness to perform behavioral tasks in which they are rewarded with plain water (Reinagel, 2018). This strategy has significant advantages for both animal welfare and scientific throughput. However, free access to 2% CA water led to a reduction in trial yield of around 30% in 2-hour daily training sessions (Reinagel, 2018), impeding its use in high-throughput behavioral paradigms. Moreover, it is not known if this approach would also work in mice. We here set out to test the safety and efficacy of CA water in mice, and explore further strategies that combine the benefits of freely accessible CA water with high trial yields.

Mice readily consumed CA water without adverse health effects. In adult mice, access to 2% CA water resulted in stable weights, and water intake similar to commonly used water restriction regimes. Giving mice free access to CA water on weekends did not affect trial numbers or task performance on subsequent training days. Specifically, mice with free access to CA water on weekends were motivated to perform many trials (500-1000 daily) evenly over the week, doing a decision-making task in which they earned sugar water. In a large dataset of mouse decision-making behavior (International Brain Laboratory et al., 2020), weekend regimes of traditional water restriction versus free CA water led to similar weight and learning curves. Free access to 2% CA water thus allows animals more control over their water intake, without negative effects on behavioral performance.

## Materials and Methods

Experiments were approved by ORBEA Animal Welfare Body at the Champalimaud Center for the Unknown (CCU; number 2016/005) and the Cold Spring Harbor Laboratory (CSHL) Institutional Animal Care and Use Committee (protocol 16-13-10-7, amendment approved 2018-07-28).

### Experimental design

The data shown in Figure 3 (cohort 4) are part of a large, public dataset (described in (International Brain Laboratory et al., 2020) and available at https://data.internationalbrainlab.org). All other experiments were collected separately (although in the case of cohort 3, using the same behavioral task and apparatus as described in (International Brain Laboratory et al., 2020); see also ‘Decision-making task’).

#### Cohort 1

Seventeen mice (male and female wild-type Thy1-GCaMP6, C57BL/6 background, 2-3 months) were single-housed. Each animal’s baseline weight was recorded on five consecutive days, while they had free access to plain water in their home cage. They were then divided into three groups, each with its own water regime. The first group (n = 8) received measured amounts of water each weekday. This was either 600 μL, or 40 μL of water per gram of body weight as measured right before water administration (range 600 - 1150 μL). On Saturday they received double this amount, while no water was administered on Sundays. The second group (n = 5) had access to free water in their home cage, in which CA (1% or 2%) was dissolved. The last group (n = 4) remained on free water in their home-cage, and served as a control. No animals showed signs of dehydration, or were dropped from the study.

The home-cage bottles with regular or 2% CA water were weighed daily, and the weight change used as a proxy for the volume of water consumed. To measure thirst, each animal was placed in a cage with 2 mL of plain water for 5 minutes on days 16, 24 and 29. For the animals on measured water, these 5 minutes replaced their daily measured water administration.

Mice in this study were on average 9 weeks old on the first day of the experimental intervention. Expected weight curves were computed from weighings of age-matched animals not undergoing fluid restriction, from (The Jackson Laboratory, 2015).

#### Cohort 2

Twelve mice (five females and seven males, C57BL/6, Jackson Laboratory, 6-17 months) were co-housed with their siblings. One animal was overweight at the start of the experiment (> 45 g, Body Condition Score 5), and was excluded; we thus show the data for 11 mice in Figure 1d. No animals showed signs of dehydration, or were dropped from the study. Each animal’s baseline weight was recorded on five consecutive days, while they had free access to plain water in their home cage. We then switched their home cage water to 2% CA, and recorded their daily weight. Laboratory shutdowns due to COVID-19 required us to cease this experiment, preventing a longer-term assessment of the effects of 2% CA water on body weight in older mice.

**Figure 1.**
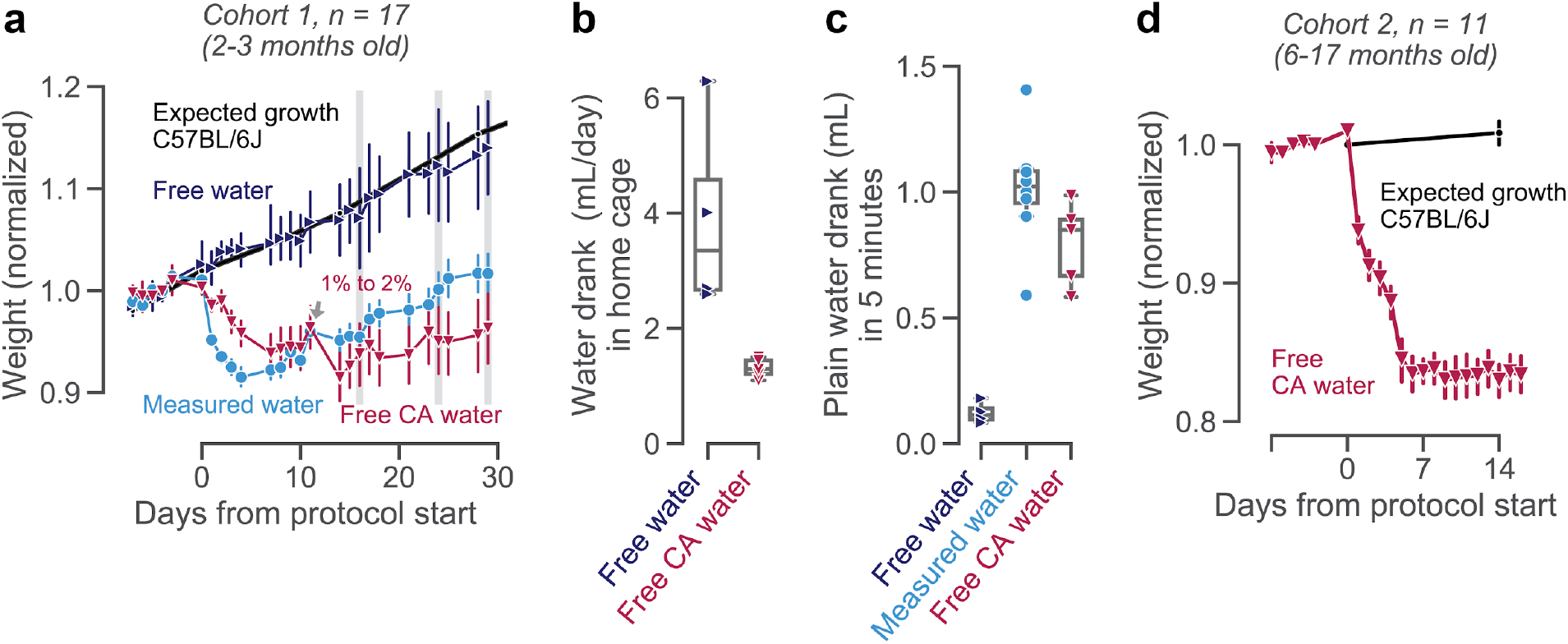
Mice maintain healthy weights and water intake on CA water. (**a**) Average weight (as a fraction of each animal’s baseline weight) for 17 young animals, divided into four experimental groups. The age-matched weight curve expected for C57BL/6J mice (The Jackson Laboratory, 2015) on free water is shown as a reference (thick black line). On day 11 (the Friday of the second week after the intervention), the CA group was switched from 1% to 2% CA in their home-cage water bottle. Error bars show mean +− 68 % confidence intervals across animals. (**b**) Volume of water drank in the home cage, estimated by measuring water bottles daily (days 11-29) for the cohorts of mice on free plain (blue) or 2% CA (red) water. Each data point is the average for one animal; boxplots show median and quartiles. (**c**) Plain water consumed in 5 minutes of free access, measured on days 16, 24 and 29, indicated with grey shaded bars in (a). Each data point is the average for one animal; boxplots show median and quartiles. (**d**) Weight curves of 11 older animals, before and after switching from free plain water to 2% CA water. Age-matched expected growth curves for C57BL/6 mice (The Jackson Laboratory, 2018) are shown as a reference in black.

Mice in this study were on average 44 weeks old on the first day of the experimental intervention. Expected weight curves were computed from weighings of age-matched animals not undergoing fluid restriction, from (The Jackson Laboratory, 2018).

#### Cohort 3

Six mice (male C57BL/6, Charles River, 8-18 months) were single-housed. On each weekday, animals performed a decision-making task (*full task,* as described below). We then varied the liquid regimes, with different types of water administration both during the weekend and weekdays (Figure 2j). Home-cage bottles were put into the cage on Friday evening and removed on Monday morning, except for the conditions where free water (regular or 2% CA) was also available on weekdays. When no home-cage bottle was available and mice did not earn their minimum required amount of 1 mL/day, they were supplemented with measured water or HydroGel at the end of the day. No animals showed signs of dehydration, or were dropped from the study.

**Figure 2.**
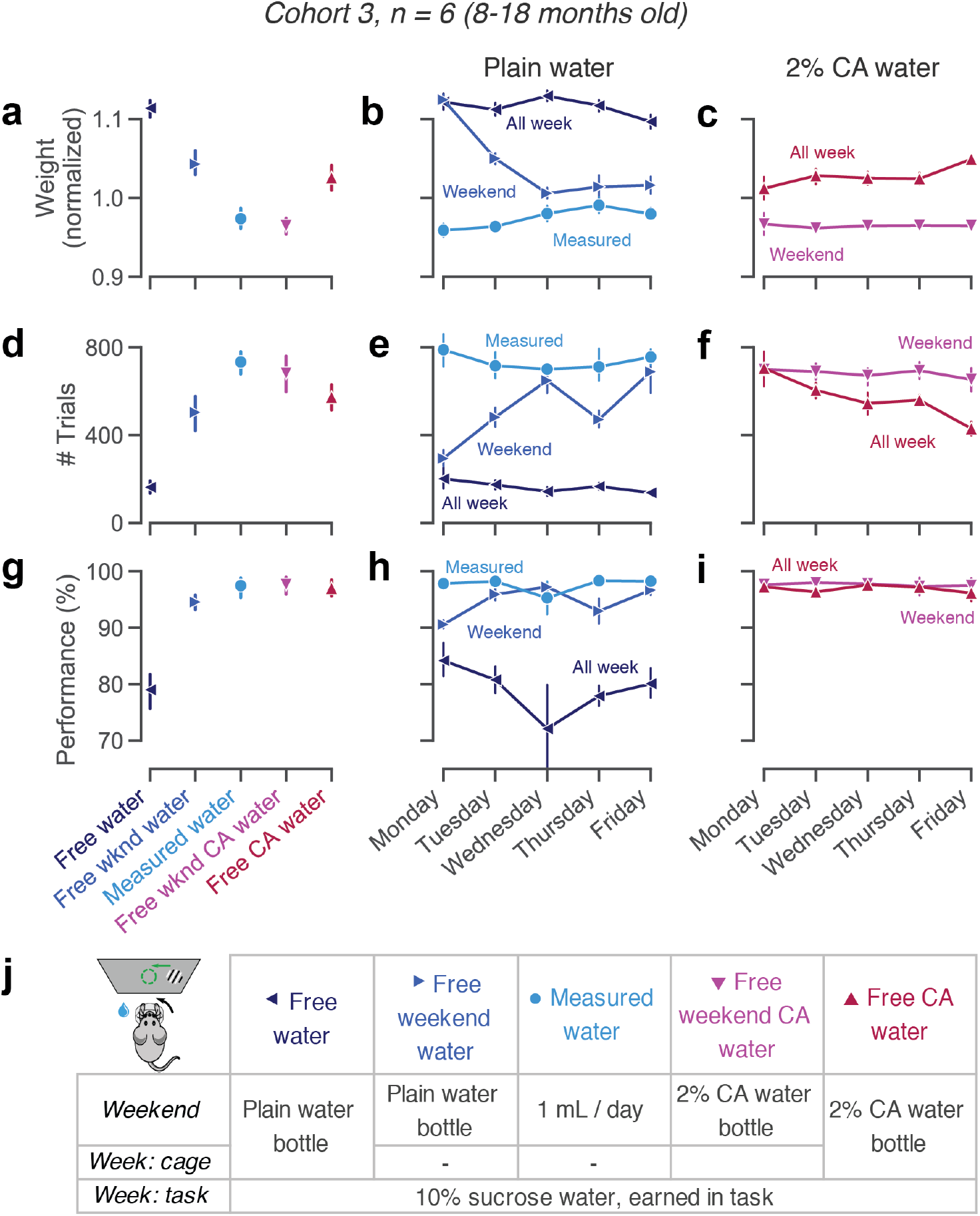
Mice perform many trials, even with free access to CA water. (**a**) Mouse weights per condition, normalized to each animal’s average weight over the course of the experiment. (**b-c**) Weights as in (**a**), per weekday. (**d**) Trials performed per condition. (**e-f**), Trials as in (**d**), per weekday. (**g**) Performance on easy trials (50% or 100% visual contrast), per condition. (**h-i**) Performance as in (**g**), per weekday. Error bars show mean +− 68% CI across animals. (**j**) Schematic of all liquid regimes. Left top corner shows a schematic of the behavioral task.

#### Cohort 4

We re-analyzed data of mice that completed training on a visual decision-making task (both *basic* and *full tasks*, see below) before 23 March 2020. All animals (140 male and female C57BL/6J, experiments performed across 7 institutions) started with a week of water restriction, handling and habituation, after which they began training. On weekdays, if they did not earn all their required water in the task, they were supplemented with HydroGel. On weekends, mice received either measured water or HydroGel (40 μL per gram of body weight or 1mL per day, depending on institutional protocols), or free access to 2% CA water. See (International Brain Laboratory et al., 2020) for detailed descriptions of the apparatus, handling and husbandry and automated training protocols. Data is available at https://data.internationalbrainlab.org.

### Decision-making task

Mice learned to perform a visual decision-making task where they detected the presence of a visual grating of variable contrast to their left or right, and reported the perceived location by turning a steering wheel (International Brain Laboratory et al., 2020). For each correct response, they received 1.5 μL of 10% sucrose water. Trial duration was estimated from stimulus onset to the delivery of feedback (either the water reward or a white noise indicating an error). The duration of each session was dependent on the engagement of the animal, which was determined through automated criteria (The International Brain Laboratory et al. 2020, their Figure 1e and Supp Table 3). Early in training, animals were supplemented towards the institutional minimum of 1 mL/day or 40 μL per gram of body after the experimental session. After they became proficient at the behavioral task, they usually performed enough trials to fully earn their water requirement in the rig on weekdays.

A mouse was considered proficient at this *basic task* once its behavior met a set of pre-specified criteria: > 400 trials performed in each of the last 3 sessions; accuracy > 80% correct on easy trials in each of the last 3 sessions; and on a psychometric function fit on those 3 sessions combined, a threshold of < 19, absolute bias of < 16, and lapses < 0.2 (International Brain Laboratory et al., 2020 - Appendix 2). Figure 3e-i shows behavioral data and psychometric function fits from the 3 days leading up to the animal being trained. The psychometric function has the form

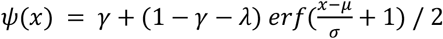

where *x* is the signed visual contrast, *μ* is a stimulus-independent bias term, *σ* is the steepness of the psychometric function, and *γ*and *λ*are lapse rates. After reaching task proficiency, animals proceeded to a more complex *full task,* where they combined visual information with an asymmetric stimulus prior that switched between blocks (International Brain Laboratory et al., 2020).

**Figure 3.**
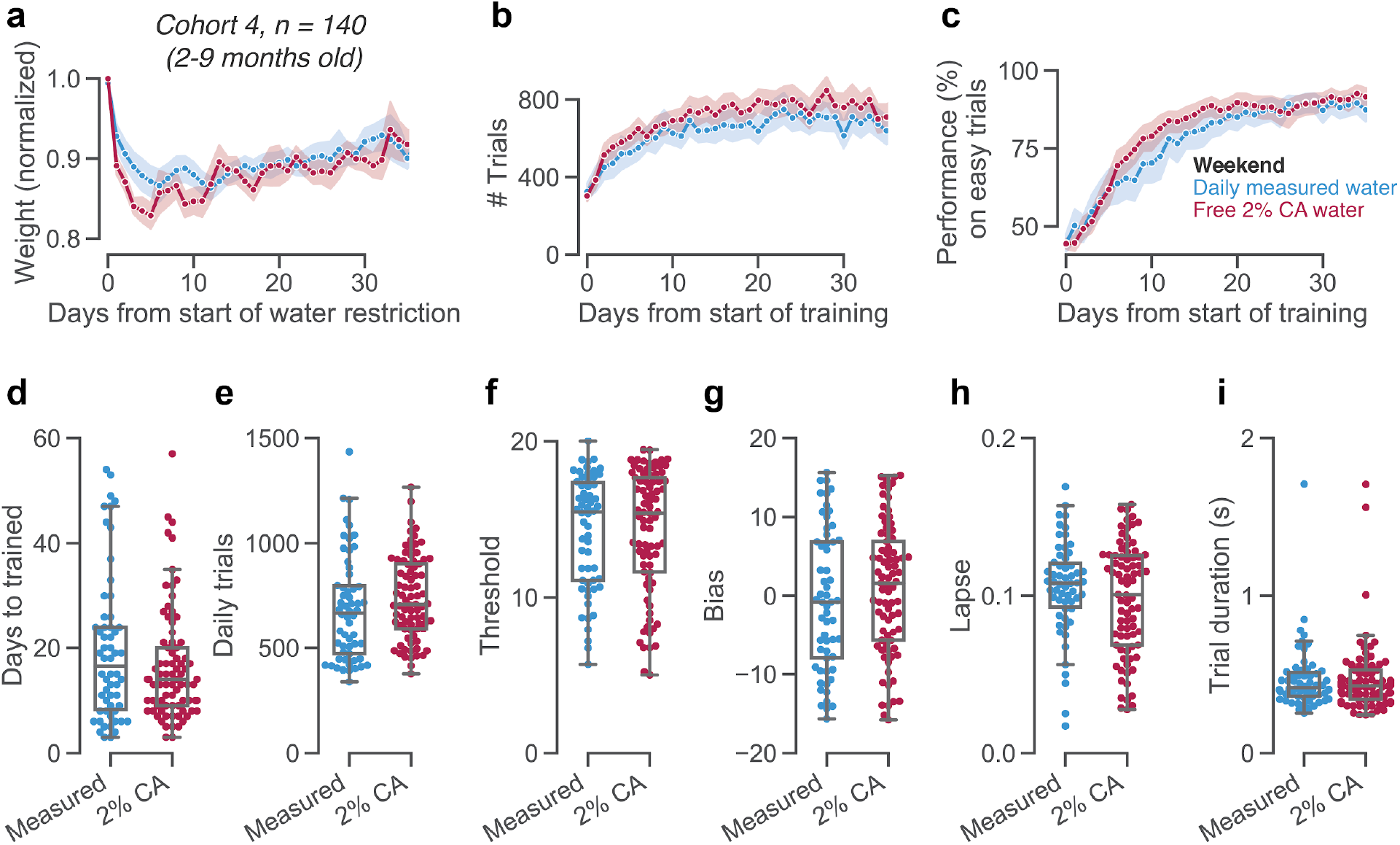
Weekend CA water does not adversely affect learning behavior. (**a**) Weight curves (normalized by each animal’s free water weight), separately for animals receiving measured water (in accordance with local IACUC protocols) or free 2% CA water on weekends and other non-training days. (**b**) Trial counts in a decision-making task over the course of learning. Training started approximately 8 days after the beginning of water restriction. (**c**) Learning curves, showing performance on easy trials with 50% or 100% visual contrast. (**d**-**i**) Various measures of learning rates and stable behavior, separated by the weekend regime used in each lab. (**d**) Number of days until reaching training criteria. (**e**-**i**) For the three days over which animals passes training criteria, (**e**) average number of trials performed per day, (**f**) threshold from a psychometric function fit, (**g**) choice bias from a psychometric function fit, (**h**) lapse rate from a psychometric function fit, and (**i**) median trial duration.

### CA water preparation

Citric acid (Sigma Aldric) was dissolved into tap water at 1% or 2% mass/volume (m/v). That is, at 2% m/v, 2g of CA powder was mixed into 100 mL water and shaken until dissolved. Bottles were replaced with fresh CA water weekly, or when empty.

### Data analysis

Data about each animal (sex, date of birth, lineage), their water intake and weights were logged in the Alyx colony management system (The International Brain Laboratory et al., 2019) and analyzed using DataJoint (Yatsenko et al., 2018). We visualized all data in python, using pandas and seaborn (Waskom et al., 2020). Statistics were done using the Pingouin package (Vallat, 2018).

For psychometric function parameters, as well as learning rates, daily trial counts and trial durations, we report statistics from an independent t-test to test the effect of weekend water regime (Figure 3). The degrees of freedom were corrected for unequal variances using a Welch–Satterthwaite correction. The corresponding Bayes factor indicates evidence for the null hypothesis of no difference, when smaller than 1.

### Code and data availability

All code and instructions for reproducing figures and statistics are available at https://github.com/int-brain-lab/citricAcid.

## Results

### Mice maintain healthy weights and water intake on CA water

We first confirmed that mice remained healthy and maintained stable weights when given free access to CA water (Figure 1a). Three subgroups of young animals (cohort 1: total n = 17, 2-3 months old) were given measured amounts of daily water, free plain water, or free CA water in their home cage. All animals were healthy (as judged by frequent experimenter handling and inspection) and showed no signs of dehydration. Over time, mice with access to free plain water steadily gained weight (Figure 1a, dark blue), as expected for young C57BL/6J mice with access to free food and water (Figure 1a, thick black line). Mice on measured water rapidly lost weight over the first week, after which their weight loss reversed (Figure 1a, light blue line), matching previous reports (Guo et al., 2014). Mice with access to CA water at a low concentration (1% m/v) did not show adverse health effects nor signs of dehydration, and lost only modest amounts of weight (Figure 1a, red line). These animals also tolerated CA at a higher concentration of 2% m/v (Figure 1a), in line with findings in rats (Reinagel, 2018). With free access to CA water, animals’ weights stabilized after about a week (Figure 1a, day 11-30).

Since these young animals (on average 9 weeks old at the start of the experiment) were still growing, their weights did not solely reflect the water regime. Taking into account this expected weight gain with age, animals on free CA water showed stable weights around 78-95% (Extended Data, Figure 1 - 1a,b). We also repeated weight measurements in older mice (Figure 1d). After switching to free CA water, animals were healthy and maintained stable weights (Figure 1d). In the second week after introducing 2% CA water, animals retained on average 83% (78-91%) of their baseline body weight (Extended Data, Figure 1 - 1c). While a weight loss of 20% is considerably larger than for rats with free access to 2% CA water (who retained 97% of their baseline body weight, Reinagel, 2018), it is similar to those stable weights obtained under widely used water restriction protocols (75-85% after several weeks of water restriction, (Guo et al., 2014)).

As expected from CA-induced weight loss, animals with access to CA water reduced their fluid intake (Figure 1b,c). Mice consumed 3.9 mL of plain water per day (range 2.6-6.3 mL; Figure 1b, blue), but only about one third as much 2% CA water (1.3 mL, range 1.3-1.5 mL; Figure 1b, red). Importantly, mice with access to free 2% CA water were still motivated to drink plain water. In five minutes of free access, they drank 0.58-0.99 mL plain water, similar to animals receiving daily measured water (Figure 1c). Not only did animals tolerate 2% CA water well, these data also suggest that free access to CA water might not impede animals’ motivation to perform behavioral tasks to earn more palatable liquids.

### Mice perform many trials, even with free access to CA water

In a new cohort of animals (cohort 2: n = 6, 8-18 months) that had been trained on a visual decision-making task (International Brain Laboratory et al., 2020), we tested how different liquid regimes affected animals’ willingness to work for sweetened water rewards (Figure 2). Animals performed a decision-making task for 45-90 minutes each weekday, and earned a drop of 10% sucrose water for each correct response. The length of each session was dictated by the automated detection of the animal’s engagement, allowing them to work to satiety (The International Brain Laboratory et al., 2020, their Figure 1e and Supp Table 3). Expert animals usually earned more than their minimum daily water requirement (determined by institutional protocols) in the task, and did not require supplemental fluids. In this cohort, we then varied the liquid regime from week to week over several months (Figure 2j, Extended Data Figure 2 - 1).

On weekends and other non-training days, mice are commonly given either daily measured water, or free access to a bottle in their home cage. When mice drank measured amounts (1 mL) of plain water on weekends (light blue circle), they performed sessions of 730-1230 trials stably over the week (Figure 2e). This practice is commonly used in laboratories that perform high-throughput mouse training. It is also the most laborious for the experimenter, who needs to provide measured amounts of water each day of the weekend. On the contrary, free home-cage water during the weekend (blue, rightward-pointing triangle) resulted in lower trial counts and worse performance on Mondays and Tuesdays (Figure 2e, 2h; 97-420 trials on Mondays vs. 668-1140 trials on Fridays; t(5) = −3.951, p = 0.0109, Bf_10_ = 6.491) (Busse et al., 2011). Under this regime, weights often fluctuated dramatically over the course of the week (Figure 2b, weight loss of around 10% of body weight or approximately 3 g between Mondays and Fridays), which may have adverse effects on animal health (Rowland, 2007).

Free access to 2% CA water over the weekend (pink, downward pointing triangle) combines the best of both approaches. Without requiring experimenter intervention on days without training, animals maintained stable weights (Figure 2c), and consistently performed many trials (Figure 2f). We could not detect a significant difference in the number of trials performed after a weekend with free 2% CA water (682 trials/day, range 535-826) vs. measured water (733 trials/day, range 634-817) (t(5) = 0.880, p = 0.4193, Bf_10_ = 0.505, Figure 2d). Similarly high trial yields were achieved by providing CA dissolved in HydroGel cups, a practical alternative in cases where bottled water is difficult to provide (Extended Data, Figure 2 - 2).

Free CA water yields the most trials when given only on weekends, but can also be kept in the home cage throughout the week. When free 2% CA was available continuously (red, upward pointing triangle), the number of trials slightly decreased from Monday to Friday (435-1,019 on Mondays vs 312-513 on Fridays; t(5) = 3.002, p = 0.0300, Bf_10_ = 3.035; Figure 2f). As a result, the average number of trials was slightly lower than when the 2% CA water bottle was removed on Monday morning (average 573 vs. 682 trials, t(5) = −2.505, p = 0.0542, Bf_10_ = 1.971, Figure 2d). While both continuous and weekend-only CA water allow for stable health and fairly high motivation, a regime with continuous 2% CA water availability may not consistently yield the high trial numbers required for some experimental purposes.

### Weekend CA water does not adversely affect learning behavior

Since mice maintained high motivation with free CA water on weekends, we implemented this strategy across several laboratories in a large-scale neuroscience collaboration (International Brain Laboratory et al., 2020). All animals underwent standardized surgeries and training protocols to learn a visual decision-making task (The International Brain Laboratory et al., 2020). Due to differences in local animal care arrangements and licenses, two different protocols were used on weekends: four labs (82 mice) used free 2% CA water, whereas the other five labs (58 mice) gave measured water or HydroGel. This allowed us to investigate the effects of CA water in a large cohort of animals, trained in different laboratories across 7 institutions.

Across all labs, and irrespective of the weekend water regime, animals successfully learned the task over the course of a few weeks (Figure 3b-c). Both water regimes also resulted in similar weight curves, with a characteristic rapid drop and subsequent slow weight gain as animals learned to earn sucrose water in the task (Figure 3a). The number of days needed to complete training (determined as reaching a specified set of behavioral criteria; The International Brain Laboratory et al. 2020, their Supp Table 2), did not vary with weekend water strategy (Figure 3d, t(101) = −1.47, p = 0.145, Bf_10_ = 0.491). Upon training completion, multiple measures of animal behavior were indistinguishable between those labs using measured water versus CA water on weekends: the number of trials performed per day (Figure 3e, t(104) = 1.33, p = 0.188, Bf_10_ = 0.410), visual threshold (Figure 3f, t(128) = 0.10, p = 0.919, Bf_10_ = 0.185), choice bias (Figure 3g, t(116) = 1.05, p = 0.297, Bf_10_ = 0.304), lapse rate (Figure 3h, t(133) = −1.36, p = 0.175, Bf_10_ = 0.429) and median trial duration (Figure 3i, t(128) = −0.03, p = 0.978, Bf_10_ = 0.184). These patterns were similar for both male and female mice (Extended Data, Figure 3 - 1). This suggests that CA water is a reliable alternative to water restriction for achieving high quality mouse behavior.

## Discussion

High-throughput, reliable behavioral training is a key requirement for the use of mice in behavioral and systems neuroscience. Here, we have shown that free access to CA water is well tolerated, and motivates mice to perform many trials of a decision-making task in which they earn sugar water. We thus consider CA water a promising alternative to water restriction for some experimental regimes.

In contrast to commonly used water restriction regimes, free access to CA water allows animals full control over the amount and timing of their water intake. Home-cage access to food and CA water also enables animals to eat and drink simultaneously, which benefits their metabolism and nutrition (Fertig and Edmonds, 1969; Toth and Gardiner, 2000). In some institutions, drinking water is acidified to reduce bacterial growth (Reinagel, 2018). Since 2% CA water has a pH of 2.07, it may convey antibacterial properties that benefit animal welfare.

Beyond extending previous experiments with CA water from rats (Watson et al., 1986; Reinagel, 2018) to mice, we here propose a new variant that maintains high trial yields. Young rats with continuous access to 2% CA water perform only around 68% of their water-restricted trial counts during daily sessions (84% for live-in behavioral testing; Reinagel, 2018). Similarly, when CA water was available throughout the week, mice performed around 22% fewer trials compared to usual conditions (no home-cage water available on testing days, measured water on weekends). Such continuous CA accessibility may thus be suitable if high trial yields are not needed, or if the baseline trial count is sufficiently high that an acceptable number of trials can be collected. As an alternative, we here propose that providing CA water only on weekends and other non-training days balances animal welfare with ease of use and behavioral throughput.

While we have achieved a satisfactory balance between water intake and trial counts using a concentration of 2% CA (m/v), this concentration could be further fine-tuned based on the details of each experimental setup. Trial yield may also depend on individual animal’s taste perception (which likely varies between species and strains), task design (e.g. head-fixed vs. freely moving), and local factors such as food or environmental humidity. For instance, if only CA water is available in the home cage, sweetened task water likely further increases animals’ willingness to work. The strategy of placing animals on free CA water over the weekend may not be viable for all experiments: for example, studies investigating differences in reward and taste processing, or those requiring precise tracking of individual animal’s fluid intake.

Beyond benefits to animal welfare, eliminating water restriction also benefits the quality and throughput of behavioral experiments. It removes the need to give animals supplemental fluids on days when they do not train (e.g. weekends) or days when they do not earn their water requirements in the task (as often happens early in training). Such supplements are often beyond the capacities of routine animal care procedures, and require scientific personnel’s daily attention and record-keeping. Besides being highly labor-intensive, water supplements carry a risk of human error in the timing and precise delivery of required water amounts, as well as the distribution of water between co-housed mice. Unexpected absence of experimental personnel usually requires animals to be placed on free plain water, which can impede their progress in behavioral training for many days or weeks afterwards.

Free access to CA water provides a relatively non-labor-intensive, low-error option for keeping animals healthy without hindering behavioral training progress. This suggests that CA water can serve as a reliable and standardized strategy to achieve high quality task behavior, further facilitating the use of mice in high-throughput behavioral studies.

## Acknowledgements

We thank all members of the International Brain Laboratory behavior working group for support and advice, and Peter Dayan, Matteo Carandini, Sonja Hofer, Nick Steinmetz and Hannah Bayer for valuable comments on the manuscript. Shan Shen provided support with DataJoint queries, and Alexander Hastava and Anup Khanal assisted with the preparation of CA Hydrogels and mouse weighing.

AEU is supported by the German National Academy of Sciences Leopoldina. FC was supported by an EMBO long-term fellowship and an AXA postdoctoral fellowship. The International Brain Laboratory is supported by the Simons Collaboration on the Global Brain and the Wellcome Trust (grant numbers 209558 and 216324).

## Author Contributions

AEU, VAR, ICL and FC designed and performed research; AEU and ICL analyzed data; AEU wrote the first draft; AEU, VAR, ICL, FC, IBL, ZM and AKC edited and approved the final manuscript; IBL, ZM and AKC provided resources and hosted the work.

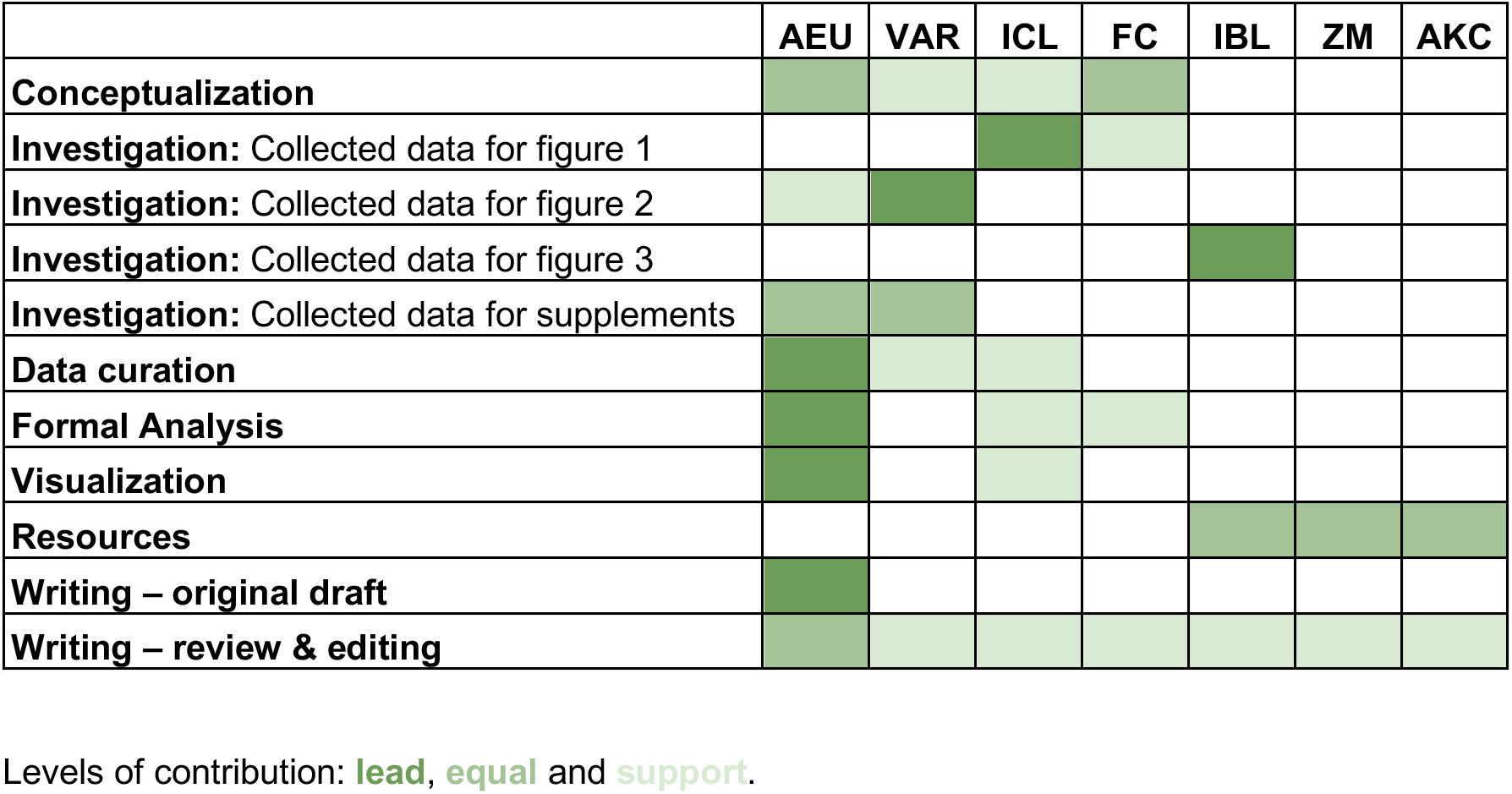

## Extended Data

**Figure 1 - 1.**
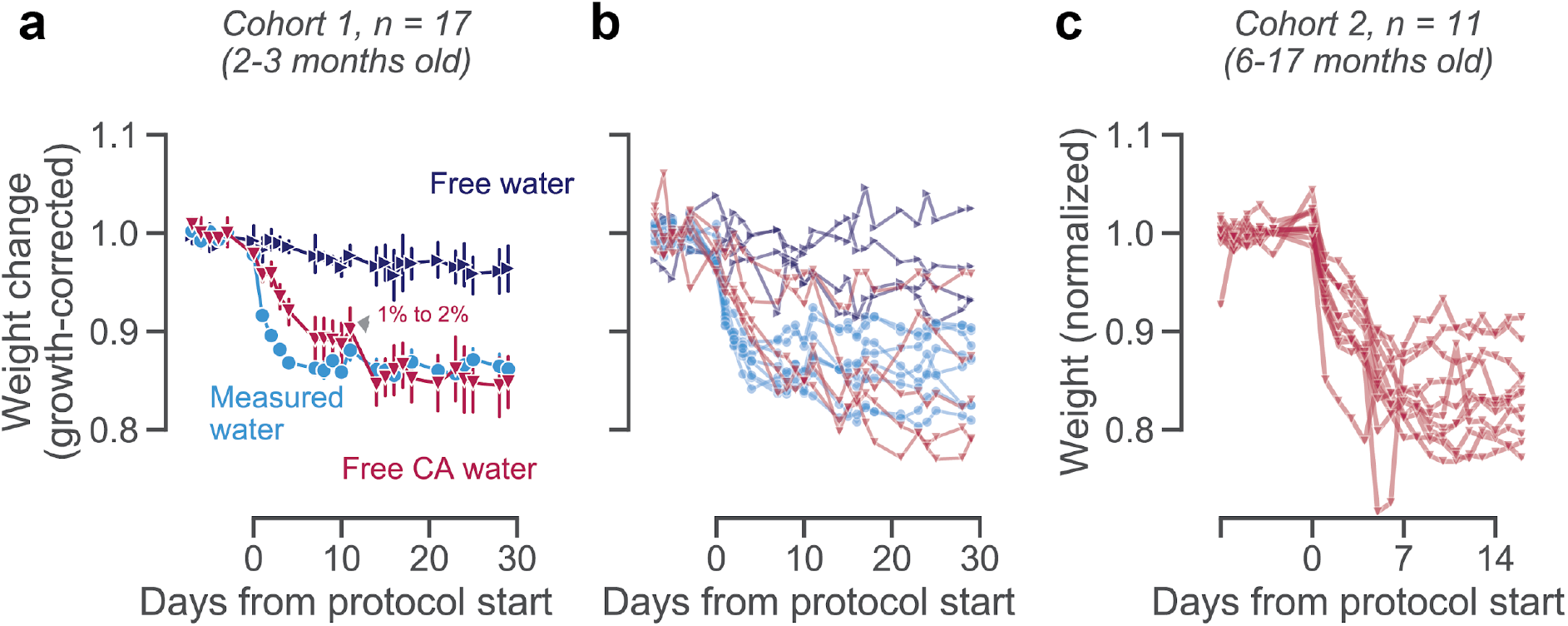
Growth-corrected weight curves. (**a**) Data as in Figure 1a, but for growth-corrected weight curves. These were computed by expressing each animal’s baseline-corrected weight as a fraction of a sex-matched expected growth curve, data from (The Jackson Laboratory, 2015). (**b**) as in a, but each animal shown individually. Young animals on CA water reached stable, growth-corrected weights of 78-95% (averaged over days 20-30). (**c**) Data as in Figure 1d, for each animal shown individually. Adult animals on CA water reached stable weights of 78-91% (averaged over days 7-14).

**Figure 2 - 1.**
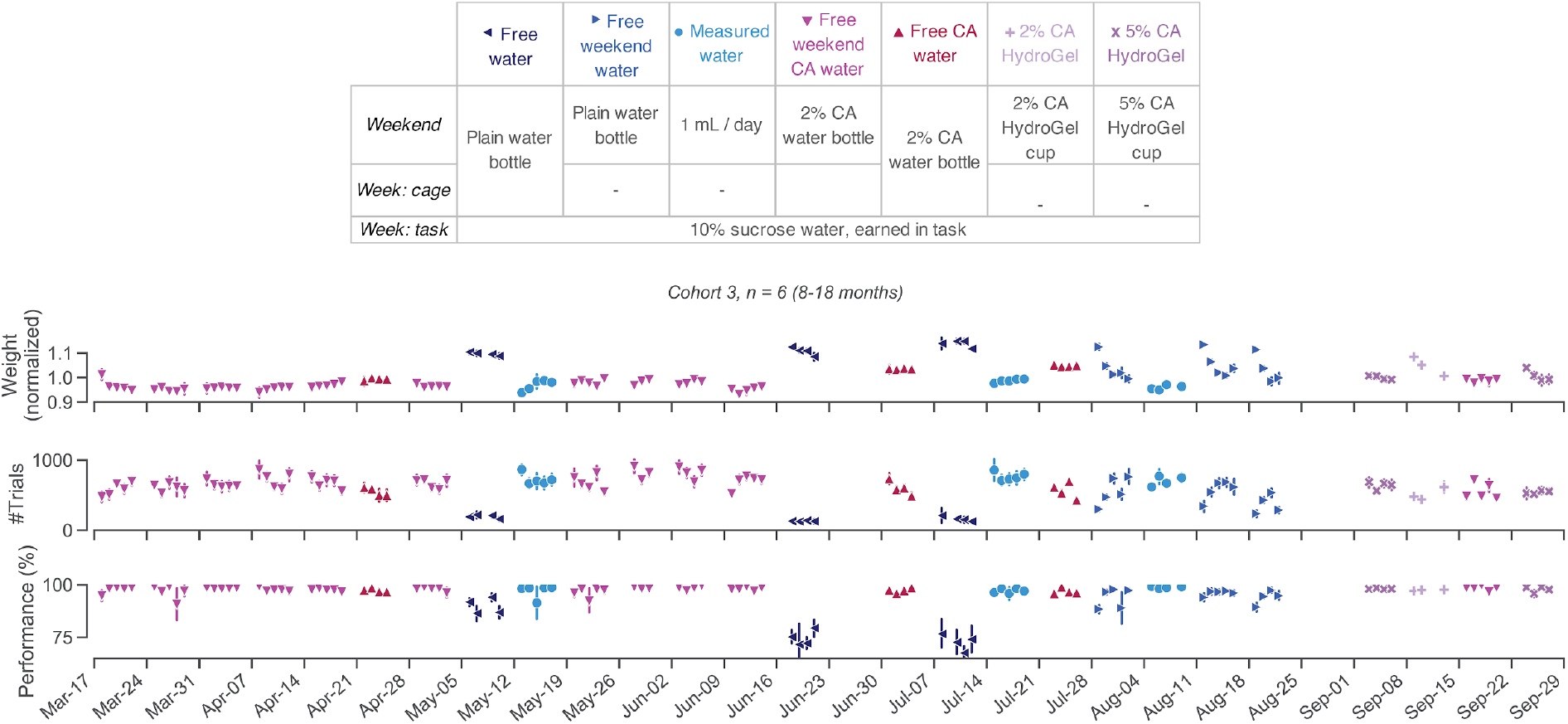
Weight, trial counts and behavioral performance over time. Data as in Figure 2, shown over the full period of data collection. A leak in the rig tubing resulted in inaccurate reward volumes during two weeks of training (in late June and early September); these data were excluded from all analyses.

### Figure 2 - 2. Citric acid can be dissolved in Hydrogel ™ instead of water

If bottled CA water cannot easily be provided (during travel, due to cage size restrictions, or when head implants preclude the use of bottle-top cages), CA can be mixed into HydroGel− cups (clearh2o.com/product/hydrogel) as an alternative to liquid water. HydroGel− was melted by placing unopened 56g cups in a 60°C oven until the gel had liquified (3-5 hours). CA powder was then mixed into the liquefied gel and stirred thoroughly, before resealing the HydroGel cups and letting them solidify at 4°C. As the flavor and perceived aversiveness of CA may differ when dissolved in water or HydroGel, we again titrated CA concentrations to achieve stable animal weights. The observation that higher concentrations of CA are required in Hydrogel to achieve the same behavioral effects agreed with informal human flavour perception of both substances.

In a first cohort of animals (cohort 5: n = 5, 15 months), this required increasing concentrations from 1% to 6% m/v (Figure 2 - 2a). With a second cohort of animals (cohort 6: n = 10, 15-16 months), switching from plain Hydrogel to 6% CA Hydrogel resulted in weights close to the institutional minimum of 80%. At 5% CA Hydrogel, all animals showed stable weights (Figure 2 - 2b).

In our cohort of trained animals (Figure 2), we confirmed that a weekend regime of 2% CA could be replaced by 5% CA in Hydrogel (Figure 2 - 1 and Figure 2 - 2). Specifically, neither weekly trial counts (t(5) = −0.932, p = 0.3941, Bf_10_ = 0.522) nor performance on easy trials (t(5) = −1.444, p = 0.2083, Bf_10_ = 0.771) were significantly different in weeks following 2% CA water vs. 5% CA HydroGel.

**Figure 2 - 2.**
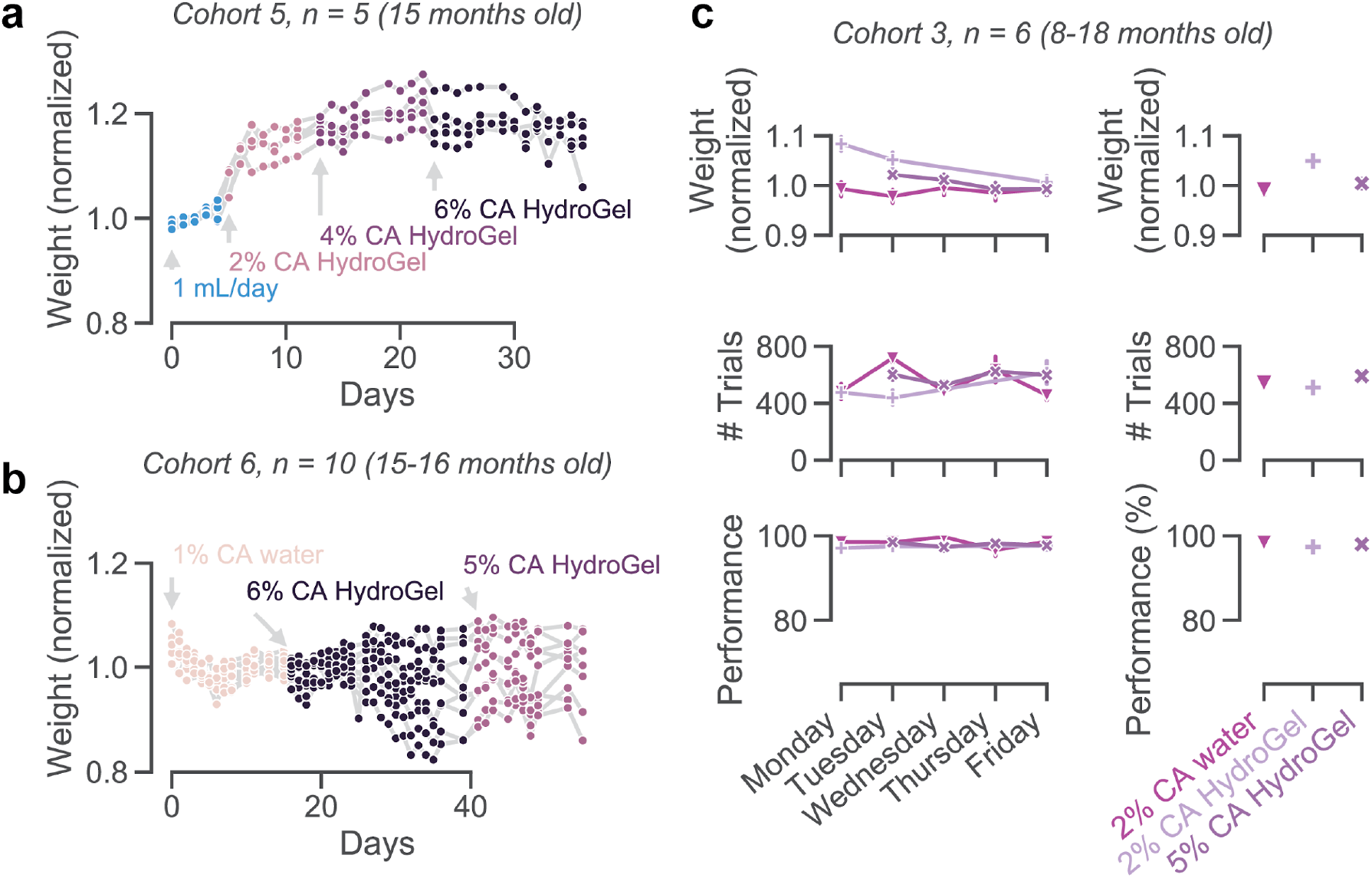
(**a**) Weight (from baseline on daily measured water), as animals were given free HydroGel with different concentrations of CA. (**b**) As in (**a**), but with a larger cohort. (**c**) As in Figure 2, but comparing 2% CA water with 2% CA HydroGel and 5% CA HydroGel on weekends (same animals as shown in Figure 2; see also Figure 2 - 1).

### Figure 3 - 1. Sex differences

We tested if the weekend water regime (2% CA bottle vs. measured water) differently affected male and female mice. Female mice given measured water on weekends learned the task slightly slower than female mice given 2% CA on weekends (t(27) = −2.89, p = 0.007, Bf_10_ = 7.731). This was not the case for male mice (t(77) = 0.30, p = 0.762, Bf_10_ = 0.242). Learning speeds showed a main effect of sex, and a significant interaction between water regime and sex (two-way ANOVA: effect of sex F(1) = 14.367, p < 0.001; effect of water regime F(1) = 4.645, p = 0.033; interaction F(1) = 9.457, p = 0.003). The overall slower learning speeds of female mice may be due to their lower weights, causing them to be satiated more quickly and performing fewer trials early in the training process (we gave all animals a fixed reward volume, independent of their body weight). We can speculate that animals’ weight and hydration balance may be slightly different in different water regimes, which interacts with motivation and learning speed in a sex-specific manner. Learning speeds differ between labs, which may be caused by various factors (International Brain Laboratory et al., 2020). Further work is thus needed disentangle any sex differences in the effects of water regime on task learning.

There was no significant effect of sex, or interaction between sex and water regime, for stable behavior upon training completion: daily trial counts (effect of sex F(1) = 0.345, p = 0.558; effect of water regime F(1) = 2.070, p = 0.153; interaction F(1) = 0.061, p = 0.806), visual threshold (effect of sex F(1) = 0.142, p = 0.707; effect of water regime F(1) = 0.002, p = 0.962; interaction F(1) = 0.584, p = 0.446), choice bias (effect of sex F(1) = 0.825, p = 0.365; effect of water regime F(1) = 1.374, p = 0.243; interaction F(1) = 0.027, p = 0.869), lapse rate (effect of sex F(1) = 2.085, p = 0.151; effect of water regime F(1) = 2.284, p = 0.133; interaction F(1) = 0.320, p = 0.573) or trial duration (effect of sex F(1) = 2.730, p = 0.101; effect of water regime F(1) = 0.064, p = 0.801; interaction F(1) = 2.457, p = 0.119).

**Figure 3 - 1.**
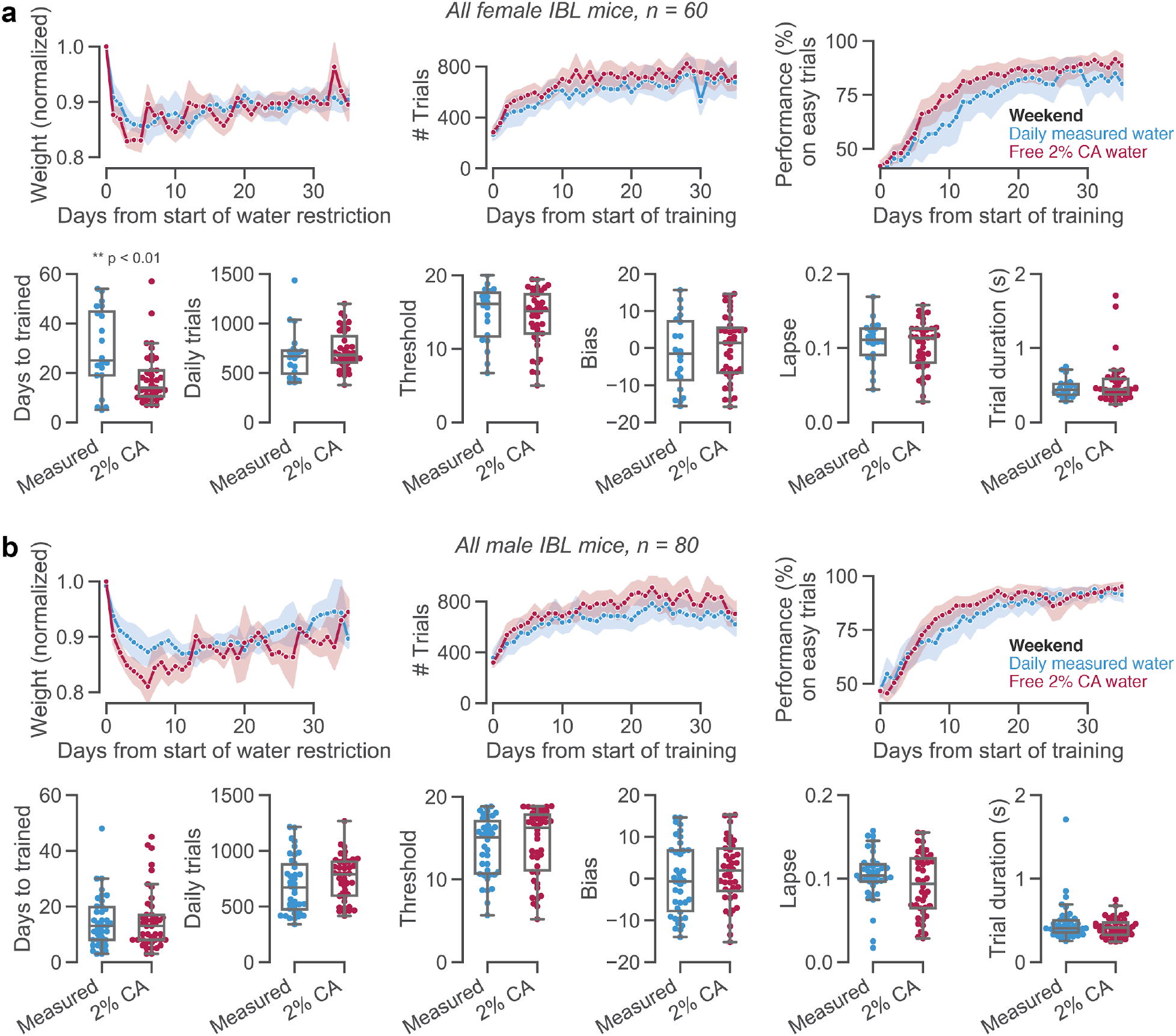
As in Figure 3, separately for (**a**) female and (**b**) male mice.

## References

Abbott A (2010) Neuroscience: The rat pack. Nat News 465:282–283.

Busse L, Ayaz A, Dhruv NT, Katzner S, Saleem AB, Schölvinck ML, Zaharia AD, Carandini M (2011) The Detection of Visual Contrast in the Behaving Mouse. J Neurosci 31:11351–11361.

Carandini M, Churchland AK (2013) Probing perceptual decisions in rodents. Nat Neurosci 16:824–831.

Fertig DS, Edmonds VW (1969) The Physiology of the House Mouse. Sci Am 221:103–113.

Goltstein PM, Reinert S, Glas A, Bonhoeffer T, Hübener M (2018) Food and water restriction lead to differential learning behaviors in a head-fixed two-choice visual discrimination task for mice. PLOS ONE 13:e0204066.

Guo ZV, Hires SA, Li N, O’Connor DH, Komiyama T, Ophir E, Huber D, Bonardi C, Morandell K, Gutnisky D, Peron S, Xu N, Cox J, Svoboda K (2014) Procedures for Behavioral Experiments in Head-Fixed Mice. PLoS ONE 9:e88678.

International Brain Laboratory et al. (2020) A standardized and reproducible method to measure decision-making in mice. bioRxiv:909838.

Reinagel P (2018) Training Rats Using Water Rewards Without Water Restriction. Front Behav Neurosci 12:84.

Rowland NE (2007) Food or fluid restriction in common laboratory animals: balancing welfare considerations with scientific inquiry. Comp Med 57:149–160.

Skinner BF (1936) Thirst as an Arbitrary Drive. J Gen Psychol 15:205–210.

The International Brain Laboratory, Bonacchi N, Chapuis G, Churchland AK, Harris KD, Rossant C, Sasaki M, Shen S, Steinmetz NA, Walker EY, Winter O, Wells M (2019) Data architecture and visualization for a large-scale neuroscience collaboration. bioRxiv:827873.

The Jackson Laboratory (2015) Body Weight Information for C57BL/6J (000664). Available at: https://www.jax.org/jax-mice-and-services/strain-data-sheet-pages/body-weight-chart-000664 [Accessed May 6, 2020].

The Jackson Laboratory (2018) Body Weight Information for Aged C57BL/6J (000664). Available at: https://www.jax.org/jax-mice-and-services/strain-data-sheet-pages/body-weight-chart-aged-b6 [Accessed May 6, 2020].

Toth LA, Gardiner TW (2000) Food and water restriction protocols: physiological and behavioral considerations. Contemp Top Lab Anim Sci 39:9–17.

Tucci V, Hardy A, Nolan PM (2006) A comparison of physiological and behavioural parameters in C57BL/6J mice undergoing food or water restriction regimes. Behav Brain Res 173:22–29.

Vallat R (2018) Pingouin: statistics in Python. J Open Source Softw 3:1026.

Waskom M et al. (2020) seaborn. Zenodo. Available at: https://zenodo.org/record/3629446.

Watson PJ, Beatey S, Wagner F, Stahl T (1986) Water adulteration with citric acid: Effects on drinking and responsivity to regulatory challenges. Physiol Behav 36:329–338.

Yatsenko D, Walker EY, Tolias AS (2018) DataJoint: A Simpler Relational Data Model. arXiv:1807.11104.

